# Modeling and Analysis of a Cell-Free Gluconate Responsive Biosensor

**DOI:** 10.1101/2023.01.10.523462

**Authors:** Abhinav Adhikari, Abhishek Murti, Anirudh M. Narayanan, Ha Eun Lim, Jeffrey D. Varner

## Abstract

Cell-free synthetic systems are composed of the parts required for transcription and translation processes in a buffered solution. Thus, unlike living cells, cell-free systems are amenable to rapid adjustment of the reaction composition and easy sampling. Further, because cellular growth and maintenance requirements are absent, all resources can go toward synthesizing the product of interest. Recent improvement in key performance metrics, such as yield, reaction duration, and portability, has increased the space of possible applications open to cell-free systems and lowered the time required to design-build-test new circuitry. One promising application area is biosensing. This study describes developing and modeling a D-gluconate biosensor circuit operating in a reconstituted cell-free system. Model parameters were estimated using time-resolved measurements of the mRNA and protein concentration with and without the addition of D-gluconate. Sensor performance was predicted using the model for D-gluconate concentrations not used in model training. The model predicted the transcription and translation kinetics and the dose response of the circuit over several orders of magnitude of D-gluconate concentration. Global sensitivity analysis of the model parameters gave detailed insight into the operation of the sensor circuit. Taken together, this study reported an in-depth, systems-level analysis of a D-gluconate biosensor circuit operating in a reconstituted cell-free system. This circuit could be used directly to estimate D-gluconate or as a subsystem in a more extensive synthetic gene expression program.

## Introduction

Synthetic biology seeks to program biological systems to perform user-specified functions. Traditionally, synthetic biology involved engineering the functionality of living cells using genetic tools; however, recent applications have focused on cell-free platforms, which have several advantages over their in vivo counterparts. For example, the absence of cellular growth and maintenance, and the cell wall barrier, facilitate the interrogation and manipulation of reaction conditions and direct the allocation of carbon and energy resources toward a product of interest. Further, cell-free platforms enable the use of linear DNA fragments instead of plasmids, which significantly reduces the time of design-build-test cycles. Cell-free systems, which have evolved to become essential research and industrial tools, have been reviewed by many authors [1–5]. Fundamentally, a cell-free system includes only the components required for transcription and translation (TXTL) processes, such as RNA polymerase (RNAP), ribosome, transcription and translation elongation factors, energy compounds, salts, amino acids, and ribonucleotides. There are two classes of cell-free systems: reconstituted and extract-based. Extract-based systems are prepared by lysing whole cells and separating the unwanted debris. In contrast, reconstituted systems, on the other hand, are well-defined and prepared using only the factors essential for protein synthesis: purified enzymes, tRNAs, ribosomes, elongation factors, amino acids, and energy molecules. Thus, reconstituted systems are an attractive platform for studying biomolecular interactions without unwanted interactions from other components present in extract-based systems. The unique advantages of cell-free systems over living cells make them a promising platform for applications traditionally carried out in an in vivo context, such as biosensing.

Traditionally, biosensors based on live cells have faced a significant challenge: transport limitations caused by the cell membrane. Cell-free biosensors can overcome this barrier by removing the constraint of the cell membrane and allowing direct access to the transcription and translation machinery. This leads to increased sensitivity and faster response times, essential for bio-sensing purposes. Fundamentally, there are different categories of biosensors based on their operation and utility. Transcription factor (TF) based biosensors are predominantly used to detect small molecules and ions. Transcription-factor-based biosensors utilize a transcription factor that is allosterically modified upon binding to a small molecule, thereby controlling the expression of a reporter gene. Such sensing systems have recently been used to detect water contaminants, biomarkers, and antibiotics [6–9]. Because cell-free systems can be freeze-dried and later reconstituted with water, they can be used for on-demand applications, such as [9]. Moreover, existing TFs can detect other compounds through metabolic engineering. For example, Voyvodic and coworkers [10] used the TF BenR to detect benzoic acid and then used the metabolic enzymes HipO and CocE, which convert hippuric acid and cocaine into benzoic acid, thereby expanding the range of detectable molecules.

In this study, we designed a D-gluconate responsive synthetic sensor circuit, based on the transcription repressor GntR [11–15], in the reconstituted cell-free system, PUR-Express. The sensing element of the circuit was GntR, while the readout was the fluorescence of the reporter protein, Venus. Using experimental measurements from the D-gluconate circuit, we developed a mathematical model that captured the transcription and translation dynamics of the D-gluconate sensor circuit in the PURExpress system. Further, we used this model to compute the global sensitivities of all the model parameters. Finally, we validated the model with experimental data not used for model training. This work is important because it estimated important kinetic and thermodynamic parameters and allowed a systems-level understanding of the interactions between the different biological parts in the sensor circuit.

## Materials and Methods

### Construction of the gluconate-responsive transcription units

Design of the transcription unit was adapted from the P70 series developed by Noireaux and coworkers [16]. A 16bp *E. coli* GntR operator (ATGTTACCCGTATCAT) was added between the -35 and -10 region of the P70 promoter. The -10 binding site of the RpoD holoenzyme was altered as a result from *GATAAT* in the original promoter to *TATCAT* in our modified P70 promoter (mP70). The gene sequence of the reporter protein (Venus) was added downstream of this modified promoter. The 5-prime untranslated region and the T500 terminator were unchanged from ref [16]. The second transcription unit had the GntR repressor gene downstream of the original (P70) promoter (without the GntR operator). The full constructs were ordered as linear DNA fragments, with 150-200 bp flanker sequences on both ends, from Twist Bioscience. The sequences of the GntR operator and the repressor were obtained from the literature [13, 17].

### Cell-free protein synthesis reactions

The cell-free protein synthesis (CFPS) reactions were carried out using the PURExpress In Vitro Protein Synthesis Kit (New England Biolabs Inc) in 384-well plates (Thermofisher NUNC, flat-bottom) in Varioskan Lux plate reader at 37^*°*^C. The working volume of all the reactions was 25 *µ*L, composed of the PURE solutions A (10 *µ*L) and B (7.5 *µ*L), RNase inhibitor, Murine (0.5 *µ*L), NEB E.coli RNA Polymerase holoenzyme (2 *µ*L), the linear DNA: Venus (7 nM), GntR (10 nM). For the control reactions without GntR, an equal volume of nuclease-free water was used in its stead. For the de-repression reactions, D-gluconate was added while maintaining the same total volume. The *E. coli* RNA Polymerase, Holoenzyme is the core enzyme saturated with sigma factor 70 and initiates RNA synthesis from sigma 70 specific promoters. The gene constructs were ordered as linear DNA fragments, with 150-200 bp flanker sequences on both ends, from Twist Bioscience. Prior to the cell-free protein synthesis reactions, the DNA fragments were PCR amplified using Q5^®^ Hot Start High-Fidelity 2X Master Mix (New England Biolabs Inc) following manufacturer instructions and the PCR reactions were purified using PureLink™ PCR Purification Kit (Invitrogen).

### mRNA quantification

Following each CFPS run, the total RNA was extracted from 5 *µ*L of the reaction mixture using PureLink RNA Mini Kit (Thermo Fisher Scientific) and stored immediately at -80^*°*^C. The quantitative RT-PCR reactions were done using Applied Biosystems™ TaqMan™ RNA-to-CT™ 1-Step Kit. An mRNA standard curve was used to determine absolute mRNA concentrations for each of the samples. The mRNA standards were prepared as follows: separate CFPS reactions for 5 nM of plasmids (Venus and GntR) were carried out for 2 hours. Total RNA was extracted using the full reaction volume. This was followed by the removal of 16S and 23S rRNA using the MICROBExpress™ Bacterial mRNA Enrichment Kit (Life Technologies Corporation). Lastly, the MEGAclear™ Kit (Life Technologies Corporation) was used to further purify the mRNA. The mRNA concentrations were determined using the Qubit™ RNA assay kit (ThermoFisher Scientific).

### Protein Quantification

Venus fluorescence was measured using the Varioskan Lux plate reader at 513 nm (excitation) and 531 nm (emission). The fluorescence was measured with 25 *µ*L for each of the mixtures. For all measurements, at least three biological replicates were performed. A protein standard curve was used to determine the absolute protein concentrations for each of the samples. The standard curve was prepared by using Venus standards of known concentration. To prepare the standards, histidine-tagged Venus was expressed using CFPS and purified using Ni spin column (New England Bio-labs). The concentration was measured using SDS-PAGE on Mini-PROTEAN Tetra Cell (Bio-Rad Laboratories) using Any kD™ Mini-PROTEAN® TGX Stain-Free™ Protein Gels (Bio-Rad Laboratories). This was followed by staining the gel with Bio-Safe™ Coomassie Stain (Bio-Rad Laboratories). The stained gel was visualized using the ChemiDoc MP and then the concentrations were measured using Image Lab software.

### Formulation and solution of model equations

The model formulation was similar to the previous study of Adhikari et al., [18]. To model the circuit, the concentration for mRNA *m*_*i*_ and protein *p*_*i*_ were described using the balance equations:

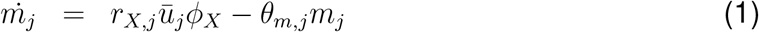

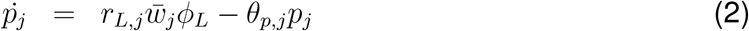

where *r*_*X,j*_ *ū*_*j*_ and 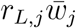 denote the transcription and translation rates, respectively (units: concentration/time), and *θ*_*m*_ and *θ*_*p*_ denote the respective degradation rate constants (units: 1/time). The *ū*_*i*_ (…) ∈ [0,1] and 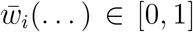 terms describe the control input governing the transcription of gene *i*, and the translation of mRNA *i*, respectively. Finally, the *ϕ*_*X*_(…) ∈ [0, 1] and *ϕ*_*L*_(…) ∈ [0, 1] terms describe the transcriptional and translational resource exhaustion, respectively.

#### Transcription and translation kinetics

The rate of transcription was modeled as the product of a kinetic limit, *r*_*X,j*_, a control term *ū*_*j*_ (…) ∈ [0,1], and the resource exhaustion term, *ϕ*_*X*_(…) ∈ [0, 1]. The kinetic limit of transcription was derived from elementary reactions (initiation, elongation, and abortive initiation) leading to the formation of the mRNA *m*_*j*_, as shown previously by Adhikari et al., [18]:

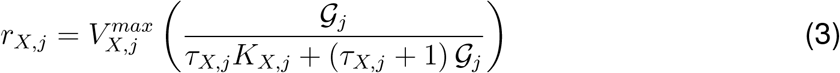

where the initiation time constant *τ*_*X,j*_, and saturation constant *K*_*X,j*_ were experimentally measured by McClure [19]. The maximum transcription rate, 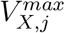, was formulated as:

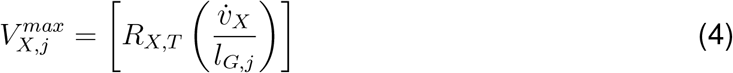

where *R*_*X,T*_ denotes the RNA polymerase concentration (nM), 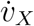 denotes the RNA polymerase elongation rate (nt/h) and *l*_*G,j*_ denotes the length of gene j in nucleotides (nt). By analogy, the kinetic limit of translation was formulated as:

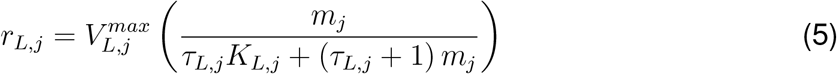

where *m*_*j*_ denotes the mRNA concentration and 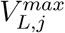 denotes the maximum translation rate formulated as:

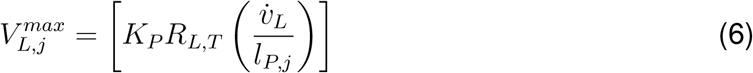

#### Transcription and translation control functions

The formulation of the control function *ū*(…) was similar to the previous study of Adhikari et al. [18] and the earlier work of Moon et al. [20]. Suppose the promoter *P* can exist in one of *s* = 1, 2 …, 𝒮 possible discrete independent microstates, where each microstate *s* has some pseudo energy *ϵ*_*s*_. Some microstates will lead to transcription, while others will not. For each microstate *s*, let’s assign a pseudo energy *ϵ*_*s*_, where *ϵ*_1_ = 0; we assume the base state of a promoter *P*, i.e., the state in which nothing is bound to the promoter-DNA has zero pseudo energy. Further, suppose the probability that promoter *P* is in microstate *s* follows a Boltzmann distribution:

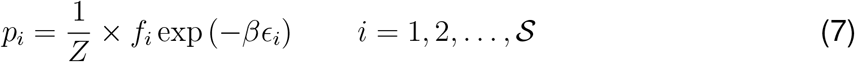

The quantity *p*_*i*_ denotes the probability that promoter *P* is in microstate *i* = 1, 2, …, 𝒮, *f*_*i*_ denotes a state-specific factor *f*_*i*_ ∈ [0, 1], *β* denotes the thermodynamic beta, and *Z* denotes a normalization factor (called the Partition function in the statistical physics community). The state-specific factors *f*_*i*_ ∈ [0, 1] were typically hill-type expressions that described the fraction of bound effector in a given microstate state; this formulation is consistent with the earlier work of Moon et al. [20].

We find *Z* using the summation law of discrete probability e.g., Σ_*s*_ *p*_*s*_ = 1 which gives:

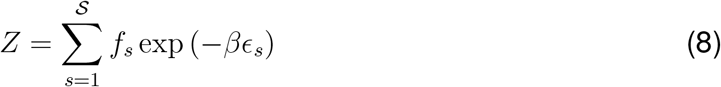

or:

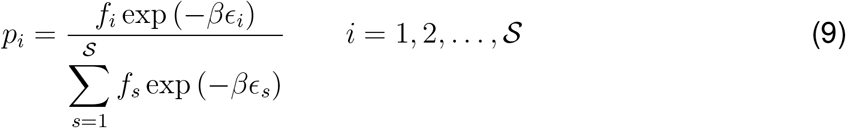

Finally, we relate the probability that promoter *P* is in microstate *s* to the *ū*(…) control function by computing the overall probability that the desired event happens, e.g., promoter *P* undergoes transcription. We know if Ω = {1, 2, …, 𝒮}, then we can define the subset 𝒜 ⊆ Ω in which the desired event happens (in this case transcription). Given 𝒜, the *ū*(…) function becomes:

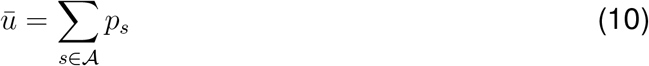

The translation control term, 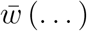, was set to 1 due to the absence of a translational regulatory network in the current system.

#### Transcription and translation resource limitation

The *ϕ*_*X*_(…) and *ϕ*_*L*_(…) functions were included in the generation terms in the mRNA (Eqn. 1) and protein (Eqn. 2) balances, respectively, to account for un-modeled resource consumption over the duration of the cell-free reaction. The *ϕ*_*X*_(…) and *ϕ*_*L*_(…) functions were formulated as:

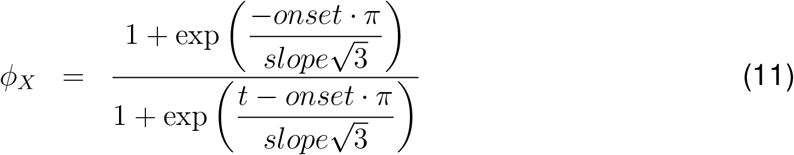

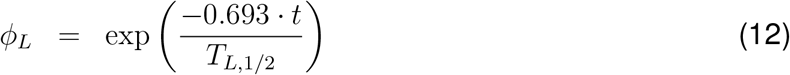

where *onset* denotes the transcription delay (measured from the start of the reaction), *slope* denotes the rate of decrease of transcription, and *T*_*L*,1*/*2_ refers to the translation half-life; values for the *T*_*L*,1*/*2_ parameter were estimated previously by Adhikari et al. [18].

The transcription decay function, *ϕ*_*X*_(…), was modeled by a logistic function that captured the slowing rate of transcription following the depletion of transcriptional resources (nucleotides, etc.) [21]. On the other hand, translation is negatively affected due to ribosome inactivation over time [22], putatively due to the build-up of inorganic phosphate, which has been shown to sequester Mg^2+^ ions [23, 24], an essential component for ribosome assembly [25, 26]. Translation decay was, therefore, modeled as a first-order decay function.

### Estimation of model parameters

Model parameter values were either taken from published studies, governed by the experimental conditions, or estimated in this study. Unknown model parameters *κ* were estimated by minimizing the squared difference between model simulations and experimental measurements generated in this study. The minimization problem was formulated as a multiobjective optimization problem, where the residual for each mRNA and protein species was treated as a separate objective, denoted as 𝒪_*j*_. The minimization problem was solved using the Pareto Optimal Ensemble Technique in the Julia programming language (JuPOETs) [27]. JuPOETs is a multi-objective optimization approach that integrates simulated annealing with Pareto optimality to estimate parameter values. JuPOETs minimized a problem of the form:

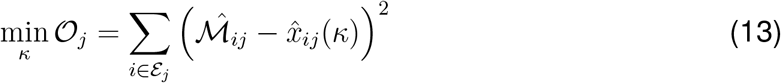

subject to:

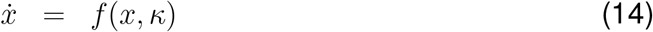

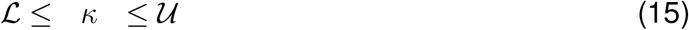

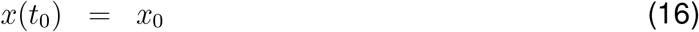

Equation (14) denotes the model equations, Eqn. (15) denotes permissible parameter bounds, and Eqn (16) represents the initial conditions. The objective function 𝒪_*j*_ quantifies the squared difference between model simulations and experimental measurements for experiment ℰ_*j*_ (measurement of either a protein or mRNA trajectory); ℳ_*ij*_ denotes the experimental value at time *i* from experiment *j*, while *x*_*ij*_(*κ*) denotes the simulated measurement at time *i* from experiment *j*; simulated *x*_*ij*_(*κ*) are implicit functions of the unknown model parameters *κ*. The lower and upper bounds for unknown model parameters *κ* were established from published studies or previous model analyses. JuPOETs was run for ten generations, all parameter ensembles with Pareto rank *≤* 5 were collected for each generation. The model equations were solved using the `DifferentialEquations.jl` package [28].

### Morris Sensitivity Analysis

The importance of model parameters was quantified using Morris sensitivity analysis, a global sensitivity analysis method [29]. The Morris method quantifies the influence of parameters on a model performance function as the mean and variance of an elementary effect. The mean of the elementary effect represents the direct influence of the parameter on system performance; a large mean value indicates a significant influence. Meanwhile, a large variance suggests that the influence of a parameter is non-linear or results from interactions with other parameters. The Morris sensitivity coefficients were computed using the `GlobalSensitivity.jl` package [30].

## Results and Discussion

### Circuit architecture and components

The genetic circuit was composed of two genes, P70-GntR and mP70-Venus (Fig. 1). The promoters for each gene were derived from the P70 series developed by Noireaux and coworkers [16]; thus, expression from both promoters was responsive to the bacterial sigma factor 70 (*σ*_70_). However, the mP70 promoter contained an additional GntR operator site between the -35 and -10 regions. The transcription factor GntR is an allosteric repressor responsive to D-Gluconate. Numerous studies on bacterial regulators, including GntR, TetR, and ArgR, have shown that these protein families have a conserved N-terminal region that promotes binding to specific operator regions on the DNA via a helix-turn-helix (HTH) motif [31–34]. On the other hand, the C-terminal sequence varies for each member in the subfamily, giving rise to different ligand binding specificities. Studies on the gluconate operon of *B. subtilis, E. coli* and *P. aeruginosa* have shown that D-gluconate acts as a de-repressor, relieving the repression of the GntR protein [11–15]. Finally, the fluorescent protein Venus was used as the reporter protein because it was previously shown to have good fluorescence properties in PURExpress [35]. All genes were encoded as PCR-amplified linear DNA, circumventing the plasmid preparation process.

**Fig. 1:**
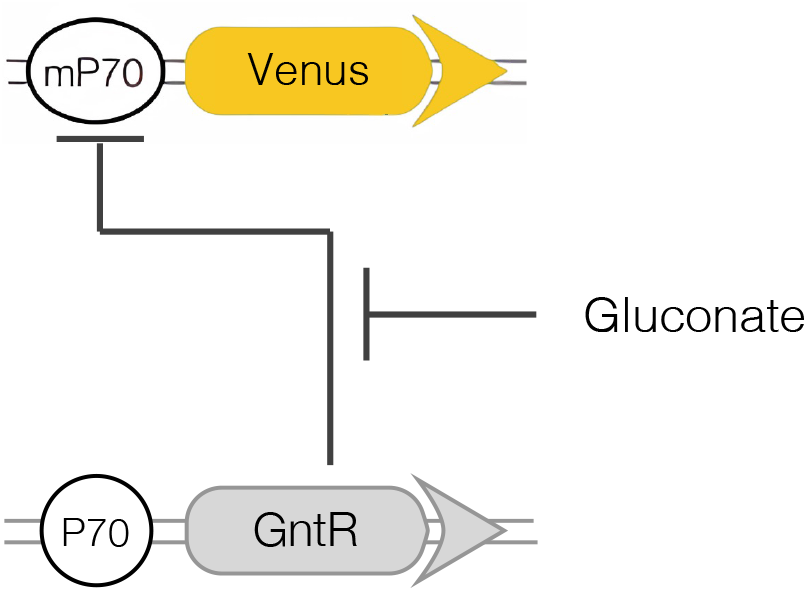
Genetic circuit architecture used in this study. The circles represent the promoters for each transcription unit. mP70 is the modified P70 promoter with the GntR operator included. The circuit components were added to the reaction as linear DNA. D-Gluconate was added as a de-repressor at the start of the reaction.

### GntR repression and de-repression by D-gluconate

The GntR protein repressed Venus expression without D-Gluconate (Fig. 2). An mP70-Venus circuit was first expressed in the reconstituted cell-free system PURExpress to characterize the yield of the fluorescent protein and its mRNA transcript. *E. coli* RNAP holoenzyme saturated with *σ*70, required for transcription, was added to the reaction mixture because PURExpress was not formulated for gene expression from an *E. coli* promoter. Transcriptional activity was maintained for 6 h, after which the mRNA began to degrade, presumably due to the combined effect of the depletion of transcription resources and the degradation of mRNA by ribonucleases (Fig. 2 A, yellow circles). On the other hand, Venus protein concentration increased slowly, and while translation continued until the end point of measurement (12 h) to yield 2.1 *µ*M, the translation rate decreased after approximately around 5 h (Fig. 2B, yellow circles). The saturating protein concentration suggested that the cell-free system’s translational capacity decreased over the reaction, which can also be attributed to the resource constraints of the cell-free extract.

**Fig. 2:**
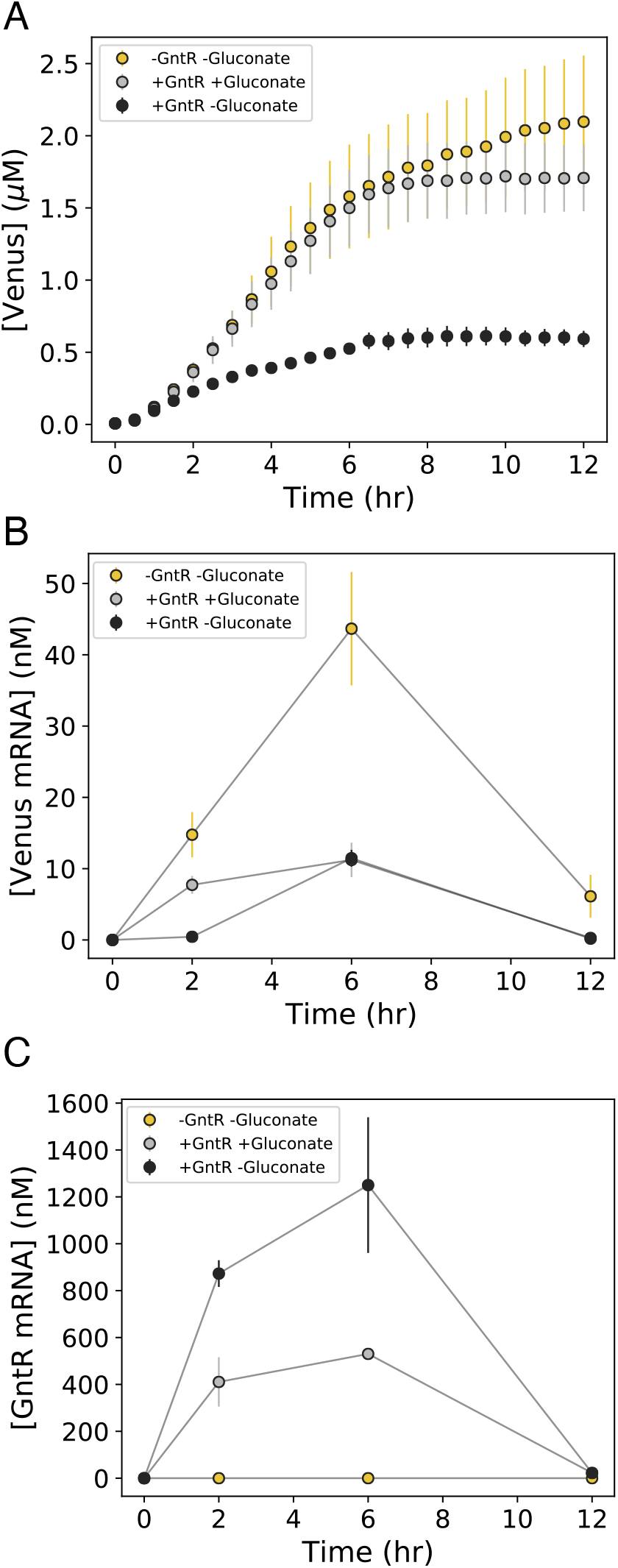
Venus mRNA and protein levels for the three different experiment conditions. Linear DNA used were: mP70-Venus (7 nM), P70-GntR (0 or 10 nM). **A**: Venus protein levels measured on a plate reader (513 nm excitation/531 nm emission). **B**: Venus mRNA levels measured using qPCR. **C**: GntR mRNA levels measured using qPCR. Yellow circles represent the unrepressed case, i.e. without the addition of GntR gene and D-gluconate. Grey circles represent the de-repressed case, i.e. with the addition of both GntR gene (10 nM) and D-gluconate (10 mM). Black circles represent the repressed case, i.e. with only the addition of GntR gene (10 nM). Error bars represent standard errors.

To characterize repression by GntR, two linear DNA fragments were expressed: *E. coli* GntR placed downstream of the P70 promoter (P70-GntR) and Venus placed downstream of the modified P70 promoter (mP70-Venus). Upon adding the P70-GntR gene, the mRNA transcript levels decreased, with the maximum Venus mRNA concentration at 6 h decreasing from 44 nM to 11 nM (Fig. 2A, black circles). Transcriptional activity was maintained for 6 h, after which the mRNA started to degrade. Venus protein expression was tightly repressed within 6.5 h (Fig. 2B, black circles); the maximum protein yield at 12 h decreased from 2.1 *µ*M to 0.6 *µ*M, a 3.5-fold change in the dynamic range. Transcript levels for GntR were more than an order of magnitude higher than those for Venus (Fig. 2C). This difference could be attributed to the change in the promoter recognition sequence in the two genes; while the -35 region in both the promoters (P70 and mP70) was unchanged, the -10 region for mP70 was changed to *TATCAT* (from *GATAAT* in P70) to accommodate the GntR operator. Additionally, to confirm if the decrease in Venus protein yield was due to repression (by GntR) and not resource limitation (owing to the addition of a second gene to the reaction mixture), the gene for a non-specific GntR variant from *P. aeruginosa* was added at the same dosage instead of the *E. coli* variant (Fig. S1). In this case, the Venus protein yield at 12 h was 1.8 *µ*M, highlighting that while the yield decreased (14%), suggesting some level of TXTL resource saturation, the repression in the previous case was indeed due to the *E. coli* GntR protein.

To study de-repression, two linear DNA fragments (P70-GntR and mP70-Venus) were expressed, with the addition of 10mM of D-gluconate (approximately 10-fold the value of *K*_*D*_ reported in literature [15]) at the start of incubation (Fig. 2, grey circles). With the addition of the effector, the Venus mRNA transcript levels at 2 h significantly increased to 7.7 nM (from 0.4 nM in the case where D-gluconate was absent), suggesting that D-gluconate prevented GntR from binding to its operator, relieving repression (Fig. 2A, grey circles). However, compared to the simple mP70-Venus circuit, Venus mRNA transcript levels were almost 2-fold lower (4-fold lower at 6 h), suggesting that repression was not entirely relieved. Despite the partial de-repression, Venus protein yield was recovered almost 80%, reaching 1.7 *µ*M at 12 h (Fig. 2B, grey circles). The GntR mRNA transcript levels were, on average, almost 2-fold lower when D-gluconate was added, suggesting that D-gluconate either lowered the transcription rate or reduced mRNA stability (Fig. 3). Taken together, GntR was functionally expressed in PURExpress; it effectively interacted with its operator and repressed transcription, and D-gluconate acted to relieve this repression. To further study de-repression, we varied the D-gluconate concentration to establish a dose-response relationship and then developed a mathematical model to further characterize this relationship in the context of several relevant kinetic and thermodynamic parameters.

**Fig. 3:**
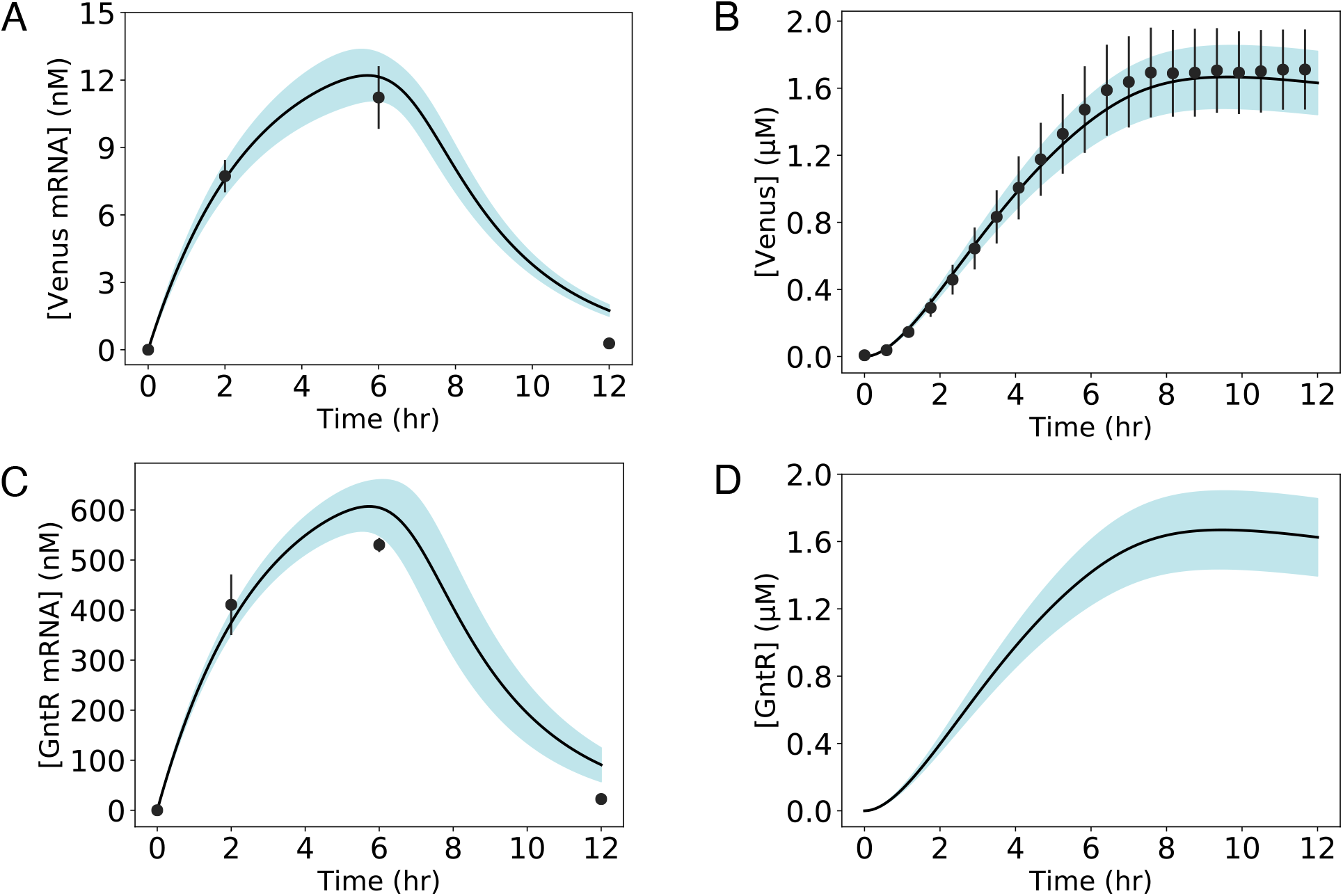
Model simulations vs. experimental measurements for the de-repressed case. Linear DNA used were: mP70-Venus (7 nM), P70-GntR (10 nM). D-gluconate (10 mM) was added as the effector at the start of reaction. **A**: Venus mRNA. **B**: Venus protein. **C**: GntR mRNA. **D**: GntR protein simulation. Black circles represent the experimental data with the standard errors captured by the error bars across at least three replicates. mRNA measurement was done using qPCR. Venus protein was measured on a plate reader (513 nm excitation/531 nm emission). JuPOETS parameter ensemble (N = 216) was used to calculate the mean (black line) and the 95% confidence estimate of the model simulation (blue region).

### Model training

We first estimated the model parameters from an mRNA and protein dataset for one experiment condition (training data set): 10 nM P70-GntR, 7 nM mP70-Venus and 10 mM D-gluconate, fixed these model parameters and then varied the effector (D-gluconate) concentration to test the predictive power of the model. The model captured *σ*_70_ induced Venus and GntR expression within the limits of experimental error for the training data set (Fig. 3). Because GntR protein was not experimentally measured, its expression was set to mirror the dynamics of Venus to ensure realistic values for its yield. JuPOETs produced an ensemble (N = 216) of the 26 unknown model parameters, which captured the expression dynamics across 12 h. The means and standard deviations of key parameters are listed in Table 1. Despite the difference in over an order of magnitude between the Venus and GntR mRNA levels, the model effectively fits both measurements. The average mRNA half-life was estimated to be 108 min, aligning closely with the value previously reported for PURExpress [36]. The decrease in transcriptional resources was modeled using a logistic distance decay function [21], added as a regulator to the transcription control function, *u*(…); the onset of decay was estimated to be 6.7 h. Venus protein expression was captured well by the model. Venus protein accumulated steadily, and its levels began to plateau after 8 h. The decrease in translational capacity was estimated using a monotonically decreasing translation capacity state variable *ϵ* and the translational control variable *w*(…). In particular, the mean half-life of translational capacity was estimated to be 3.1 h, and the average half-life for both expressed proteins was estimated to be 1.4 days.

**Table 1:** Estimated transcription and translation model parameters for the D-gluconate sensor circuit.

| Description | Parameter | Value | Units | Reference |
| --- | --- | --- | --- | --- |
| RNA Polymerase concentration | $R_X$ | 0.13 | $\mu\text{M}$ | set by experiment |
| Ribosome concentration | $R_L$ | 2.0 | $\mu\text{M}$ | set by experiment |
| Transcription elongation rate | $\dot{v}_X$ | 12-30 | nt/s | [38, 39] |
| Translation elongation rate | $\dot{v}_L$ | 1-2 | aa/s | [38, 40] |
| Transcription saturation coefficient | $K_X$ | 0.036 | $\mu\text{M}$ | [19] |
| Transcription initiation time | $k_{init,X}$ | 22 | s | [19] |
| Translation initiation time | $k_{init,L}$ | 1.5 | s | [18] |
| Venus gene length | $l_G$ | 720 | bp | from DNA sequence |
| GntR gene length | $l_P$ | 996 | bp | from DNA sequence |
| Translation saturation coefficient | $K_L$ | $76.25 \pm 2.91$ | $\mu\text{M}$ | - |
| Half-life translation | $\tau_{L,1/2}$ | $3.13 \pm 0.13$ | h | - |
| Transcription saturation onset | $T_{X,onset}$ | $6.67 \pm 0.20$ | h | - |
| Transcription saturation slope | $T_{X,slope}$ | $0.89 \pm 0.02$ | 1/h | - |
| GntR transcription | $\tau_{X,GntR}$ | $4.98 \pm 0.09$ | - | - |
| GntR translation | $\tau_{L,GntR}$ | $0.38 \pm 0.02$ | - | - |
| Venus transcription | $\tau_{X,Venus}$ | $6.14 \pm 0.32$ | - | - |
| Venus translation | $\tau_{L,Venus}$ | $0.106 \pm 0.002$ | - | - |
| half-life mRNA GntR | $\ln(2)/\theta_{m,GntR}$ | $109.14 \pm 4.52$ | min | - |
| half-life mRNA Venus | $\ln(2)/\theta_{m,Venus}$ | $106.98 \pm 4.32$ | min | - |
| half-life Protein GntR | $\ln(2)/\theta_{p,GntR}$ | $1.32 \pm 0.06$ | days | - |
| half-life Protein Venus | $\ln(2)/\theta_{p,Venus}$ | $1.49 \pm 0.07$ | days | - |
| cRNAP + P70 | $\epsilon_{P70,cRNAP}$ | $0.79 \pm 0.03$ | $k_B T$ | - |
| hRNAP + P70 | $\epsilon_{P70,hRNAP}$ | $-3.68 \pm 0.10$ | $k_B T$ | - |
| cRNAP + mP70 | $\epsilon_{mP70,cRNAP}$ | $4.87 \pm 0.06$ | $k_B T$ | - |
| hRNAP + mP70 | $\epsilon_{mP70,hRNAP}$ | $-1.48 \pm 0.07$ | $k_B T$ | - |
| GntR + mP70 operator | $\epsilon_{mP70,GntR}$ | $(-3.77 \pm 0.20)$ | $k_B T$ | - |
| Hill coefficient P70 + hRNAP | $n_{P70,hRNAP}$ | $4.11 \pm 0.18$ | dimensionless | - |
| Dissociation constant P70 + hRNAP | $K_{P70,hRNAP}$ | $10.20 \pm 0.31$ | $\mu\text{M}$ | - |
| Hill coefficient mP70 + GntR | $n_{mP70,GntR}$ | $5.48 \pm 0.18$ | dimensionless | - |
| Dissociation constant mP70 + GntR | $K_{mP70,GntR}$ | $0.54 \pm 0.02$ | $\mu\text{M}$ | - |

*Note:* cRNAP and hRNAP refer to the core and sigma factor ( $\sigma_{70}$ ) saturated versions of the RNA polymerase, respectively.

The promoter configuration pseudo energies were estimated in units of thermodynamic *β* = 1*/k*_*B*_*T*, where *k*_*B*_ denotes the Boltzmann constant and *T* denotes temperature. The model previously developed by Adhikari [18] captured the expression dynamics of mRNA and protein in synthetic cell-free circuits; however, the promoter pseudo energies were not experimentally constrained during the estimation of model parameters. Since the promoter pseudo energies are essential to the weight of a particular promoter configuration, any overestimation could lead to unrealistic simulation results. Towards this issue, in this study, we used literature values reported by Bintu and coworkers to establish bounds on the pseudo energy [37]. As expected, the pseudo energies of the RNAP holoenzyme bound to the promoters (both P70 and mP70) were several-fold higher (and more negative) than those of the RNAP core with the promoters. This difference was several orders of magnitude in terms of the weight of the respective configurations. Notably, the pseudo energy of the RNAP holoenzyme with the P70 promoter was also estimated to be higher (and more negative) than that with the mP70 promoter, verifying the low mRNA levels observed for Venus compared to those for GntR.

The importance of model parameters was quantified using Morris sensitivity analysis, a global sensitivity analysis method (Fig. 4). The sensitivity measures (mean and variance) were binned into categories based on their relative magnitudes, from no influence (white) to high influence (black). The measures for most parameters were intuitive. For example, transcription delay and slope parameters, which represented the onset time and rate of depletion of transcription resources, respectively, affected the mRNA (and protein) levels for both Venus and GntR. Similarly, the translation half-life and saturation parameters strongly affected the yield of the proteins. The binding energies of the RNAP with the promoters for Venus and GntR genes strongly affected their respective mRNA and protein species. Furthermore, the most sensitive parameters were determined to be the mRNA stability modifiers for both Venus and GntR species, which influenced both the mRNA and protein expression of the individual species. These parameters had high variances because their effect was non-linear, which was expected; the maximal translation rate had a saturating dependence on the mRNA concentration. All the parameters that affected the expression of GntR also influenced the expression of Venus. This was intuitive because GntR accumulation in the system led to the repression of Venus’s expression. On the other hand, it was not apparent that Venus mRNA stability would affect GntR expression; however, this could be explained in terms of the transcription and translation resource limitation present in the system. For example, lowered Venus mRNA levels (lower mRNA stability) would increase the share of translation resources for GntR, the accumulation of which would repress Venus mRNA transcription. This would, in turn, increase the transcription resources available for GntR. In summary, Morris sensitivity emphasized the coupled nature of the transcription and translation processes. It also highlighted the degree of influence of the parameters on the expression of the different mRNA and protein species.

**Fig. 4:**
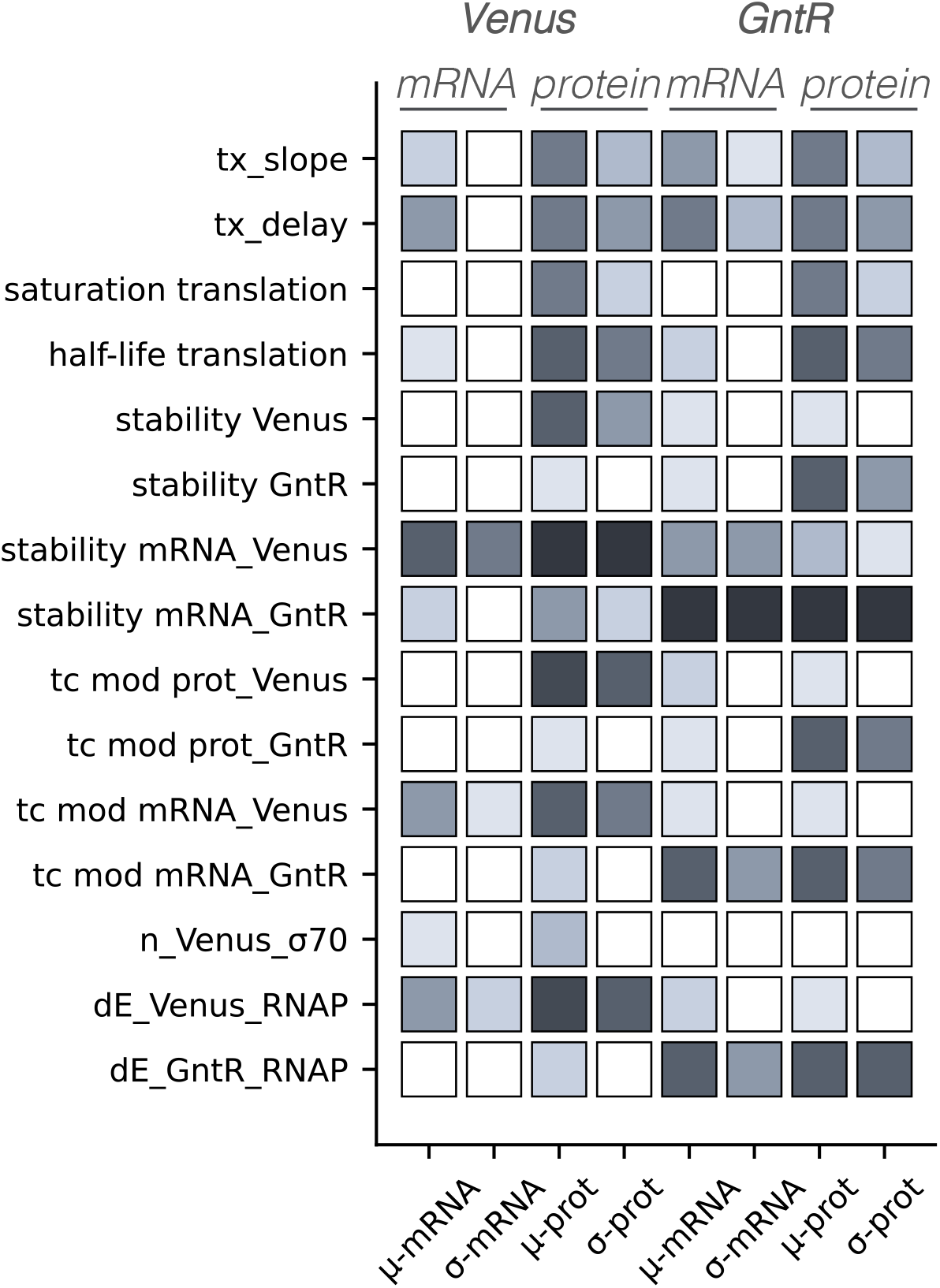
Global sensitivity analysis of the model parameters. Morris sensitivity coefficients were calculated for the estimated model parameters. *µ* represents the mean and *σ* represents the variance of the sensitivity measures. Darkness of the squares directly correlate with the sensitivity of the parameters.

#### Model prediction

In order to quantify the de-repression by D-gluconate, and test the predictive capacity of the model, a dose response study was undertaken (Fig. 5). The Venus concentration at 12 h was measured for D-gluconate concentrations ranging from 0.0001 mM to 20 mM; above 20 mM, theVenus protein expression rapidly declined (data not shown), suggesting inhibition of the PURExpress system. To simulate the dose response relationship, the parameter set fitted to the 10 mM D-gluconate case (training data set) was used. All the parameters were fixed, varying only the concentration of D-gluconate. The model captured the endpoints Venus levels within the limits of experimental error, yielding a characteristic sigmoid shape. The estimated dissociation constant *K*_*D*_ was 0.49 mM, *K*_*m*_ was 0.65 mM, and the hill coefficient (*n*) was 1.63. The model also simulated the trajectory of Venus expression over 12 h for each effector concentration (Fig. S2-S10). While the simulations closely aligned with the observed data points for most effector concentrations, it overestimated the initial trajectory for some of them. In particular, it was experimentally observed that the addition of 20 mM D-gluconate slowed down the expression of Venus (presumably due to inhibition of PURExpress); this decline was not effectively captured by the model, as evidenced by the overestimation of Venus levels upto almost 8 h.

**Fig. 5:**
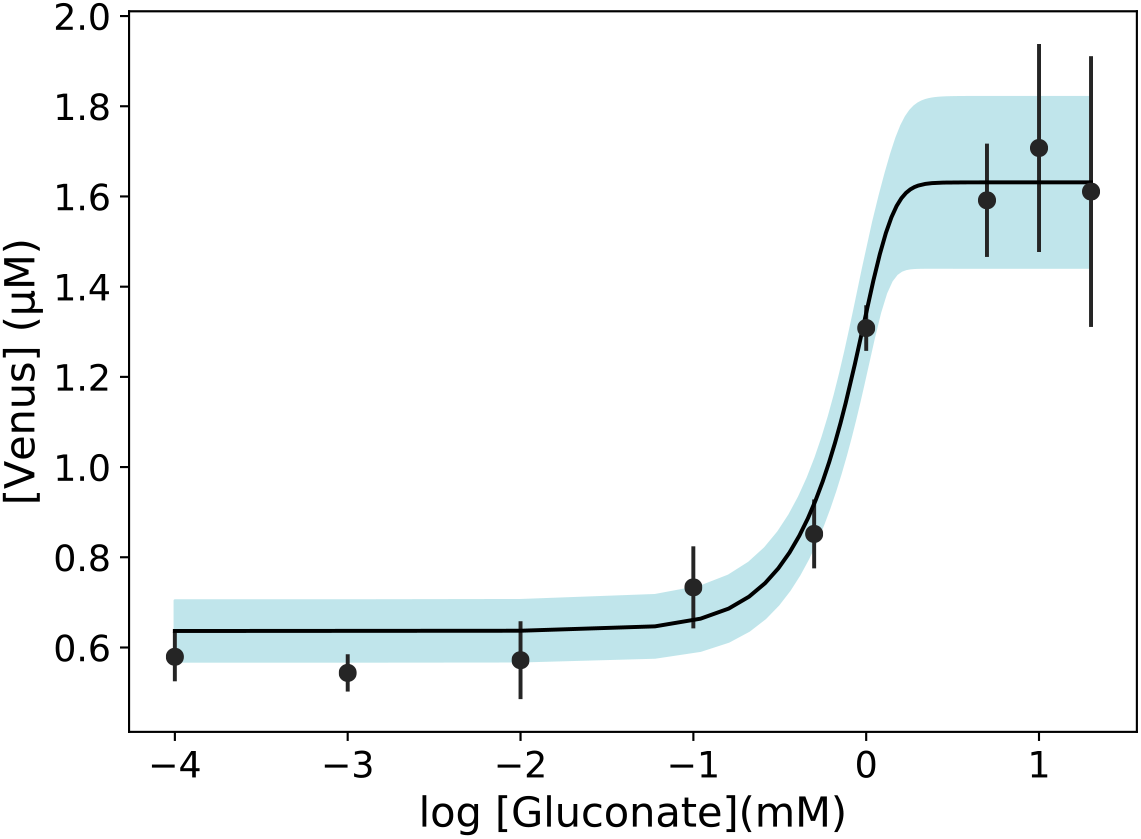
Model simulations vs. experimental measurements for the dose response study. Linear DNA used were: mP70-Venus (7 nM), P70-GntR (10 nM). D-gluconate (varied) was added as the effector at the start of reaction. Black circles represent Venus protein measurements at the endpoint of experiment (12 h). Error bars represent the standard errors across at least three replicates. Black line represents the mean of the model simulation while the blue region represents 95% confidence estimate of the model simulation.

## Author Contributions

J.V directed the study. A.A, A.N, and H.L conducted the cell free experimental measurements. A.N conducted qPCR experiments. J.V, A.M and A.A developed the reduced order models and the parameter ensemble. A.M, A.A and J.V analyzed the model ensemble, and generated figures for the manuscript. The manuscript was prepared and edited for publication by A.A, A.M, A.N and J.V. All authors reviewed this manuscript.

## Data Availability Statement

Model code is available under an MIT software license from the GitHub repository: https://github.com/varnerlab/Gluconate-biophysical-modeling-code. The mRNA and protein measurements presented in this study are available in the data directory of the model repositories in comma separated value (CSV) and Microsoft Excel format.

## Supplemental information

**Fig. S1:**
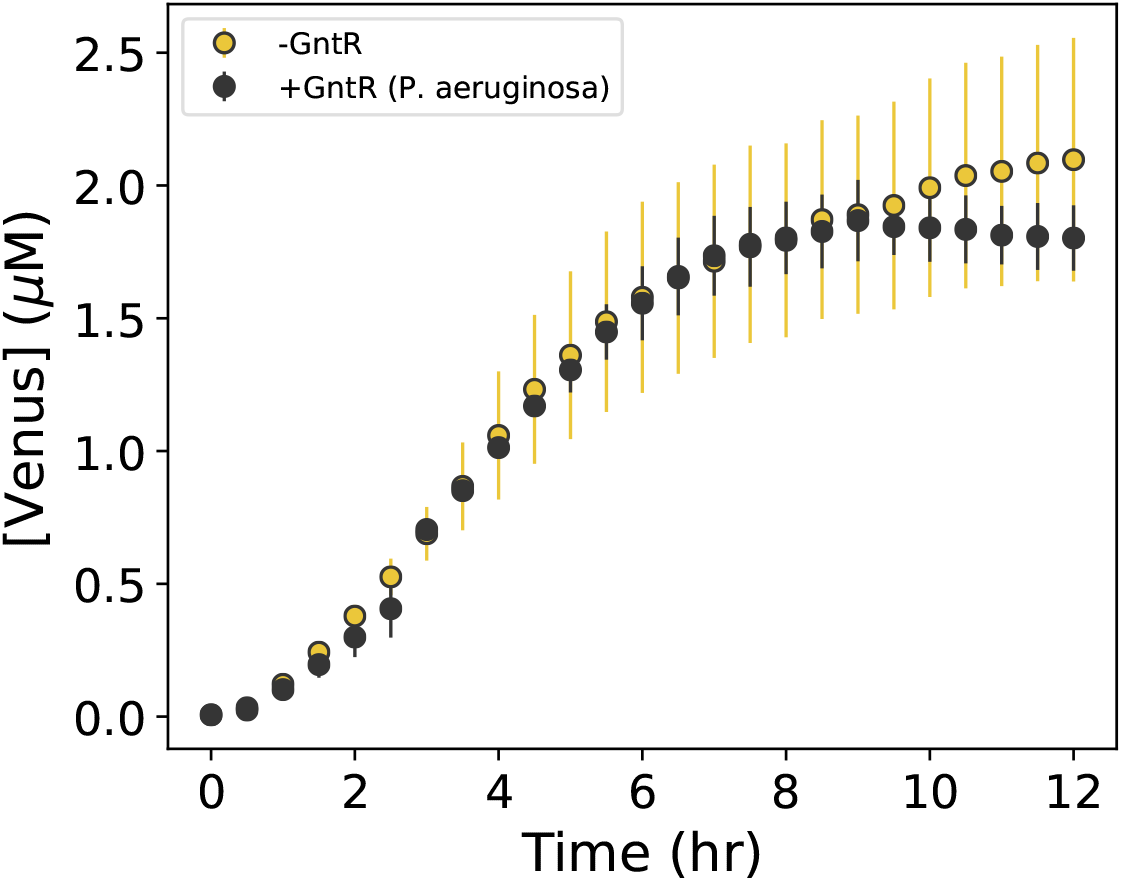
Venus protein levels measured for two cases: addition of *P. aeruginosa* GntR gene and in the absence of GntR gene. Linear DNA used were: mP70-Venus (7 nM), P70-GntR (0 or 10 nM).

**Fig. S2:**
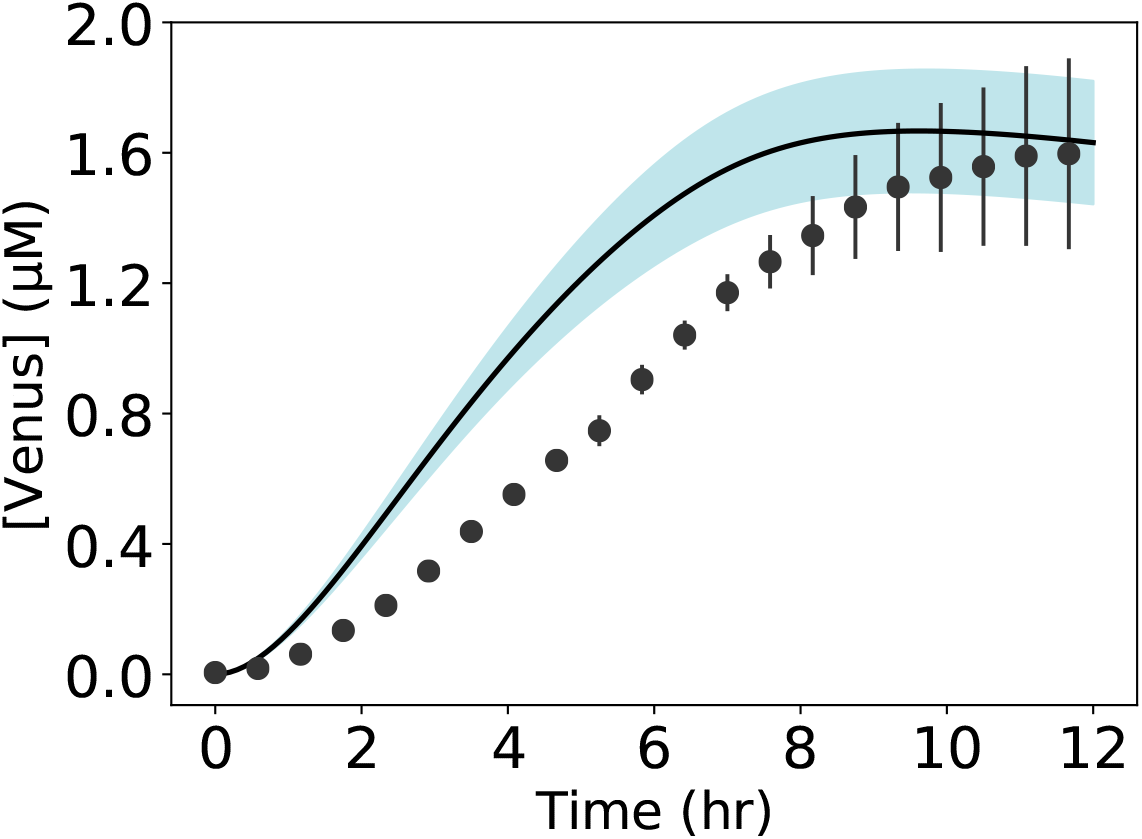
Model simulations vs. experimental measurements for the de-repressed case using 20 mM D-gluconate. Linear DNA used were: mP70-Venus (7 nM), P70-GntR (10 nM). Black circles represent the experimental data with the standard errors captured by the error bars across at least three replicates. JuPO-ETS parameter ensemble (N= 216) was used to calculate the mean (black line) and the 95% confidence estimate of the model simulation (blue region).

**Fig. S3:**
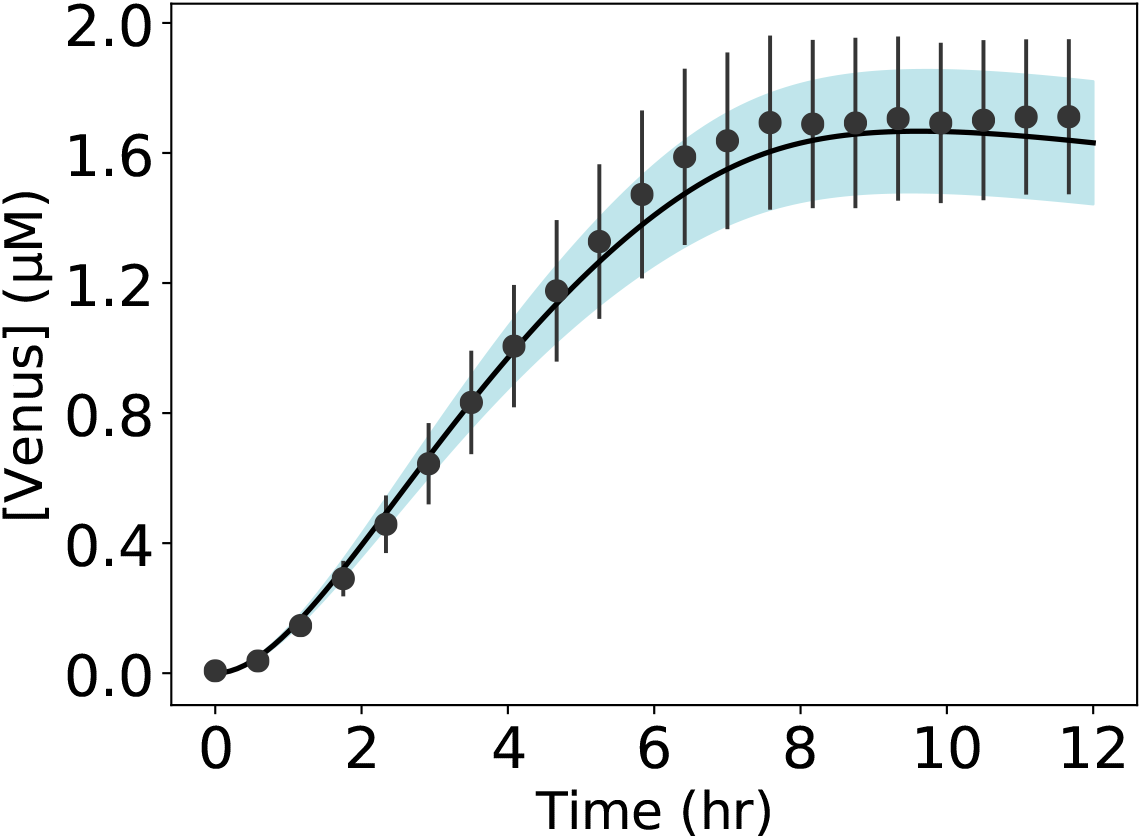
Model simulations vs. experimental measurements for the de-repressed case using 10 mM D-gluconate. Linear DNA used were: mP70-Venus (7 nM), P70-GntR (10 nM). Black circles represent the experimental data with the standard errors captured by the error bars across at least three replicates. JuPO-ETS parameter ensemble (N= 216) was used to calculate the mean (black line) and the 95% confidence estimate of the model simulation (blue region).

**Fig. S4:**
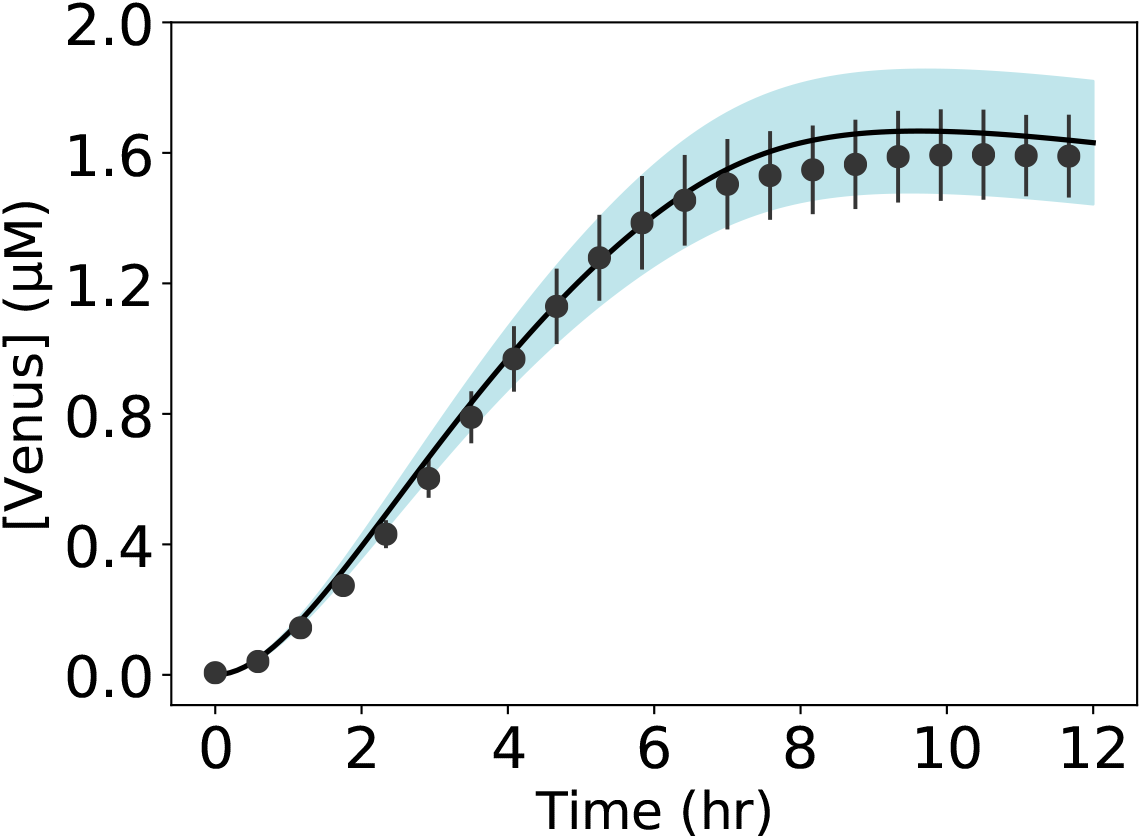
Model simulations vs. experimental measurements for the de-repressed case using 5 mM D-gluconate. Linear DNA used were: mP70-Venus (7 nM), P70-GntR (10 nM). Black circles represent the experimental data with the standard errors captured by the error bars across at least three replicates. JuPO-ETS parameter ensemble (N= 216) was used to calculate the mean (black line) and the 95% confidence estimate of the model simulation (blue region).

**Fig. S5:**
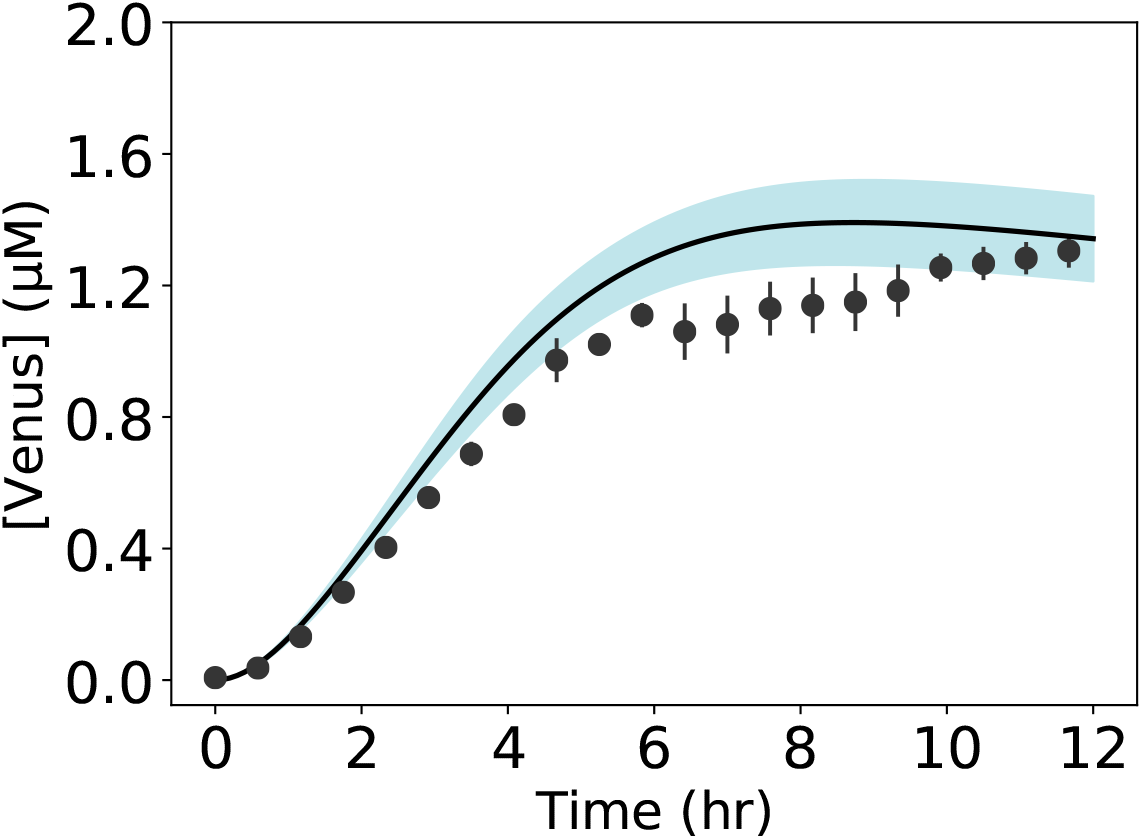
Model simulations vs. experimental measurements for the de-repressed case using 1 mM D-gluconate. Linear DNA used were: mP70-Venus (7 nM), P70-GntR (10 nM). Black circles represent the experimental data with the standard errors captured by the error bars across at least three replicates. JuPO-ETS parameter ensemble (N= 216) was used to calculate the mean (black line) and the 95% confidence estimate of the model simulation (blue region).

**Fig. S6:**
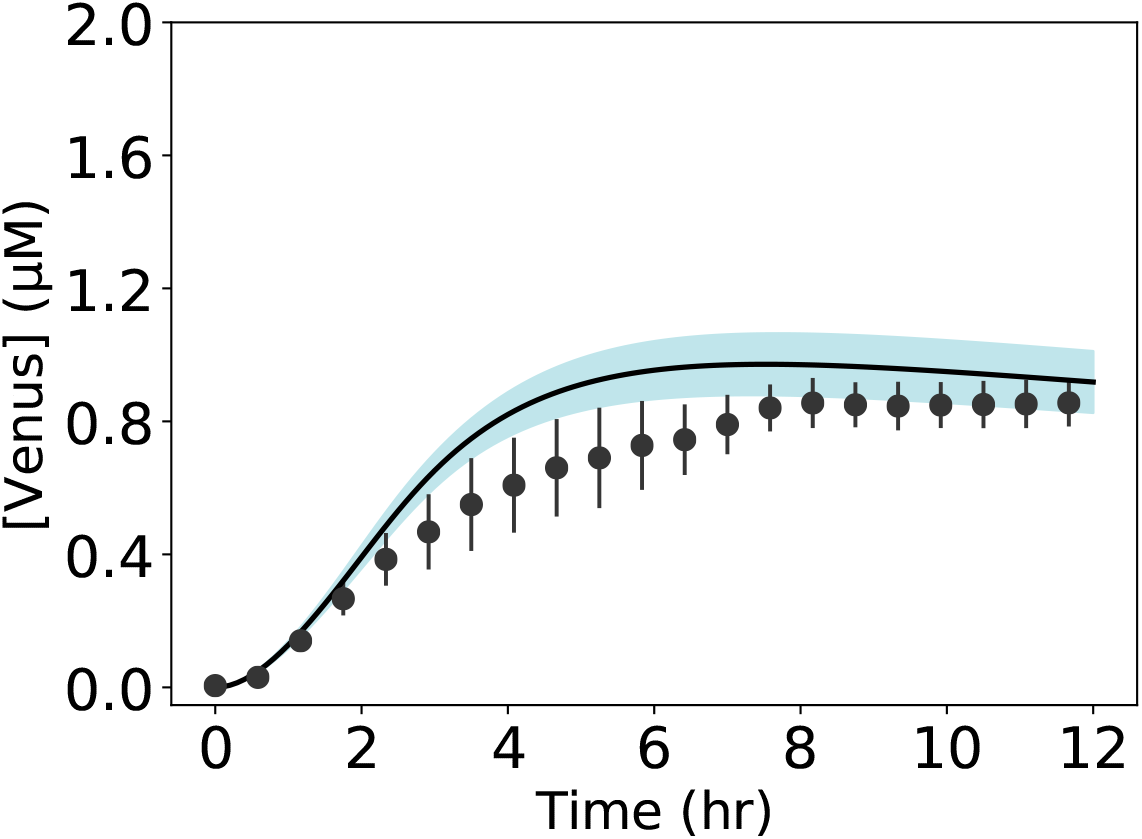
Model simulations vs. experimental measurements for the de-repressed case using 0.5 mM D-gluconate. Linear DNA used were: mP70-Venus (7 nM), P70-GntR (10 nM). Black circles represent the experimental data with the standard errors captured by the error bars across at least three replicates. JuPO-ETS parameter ensemble (N= 216) was used to calculate the mean (black line) and the 95% confidence estimate of the model simulation (blue region).

**Fig. S7:**
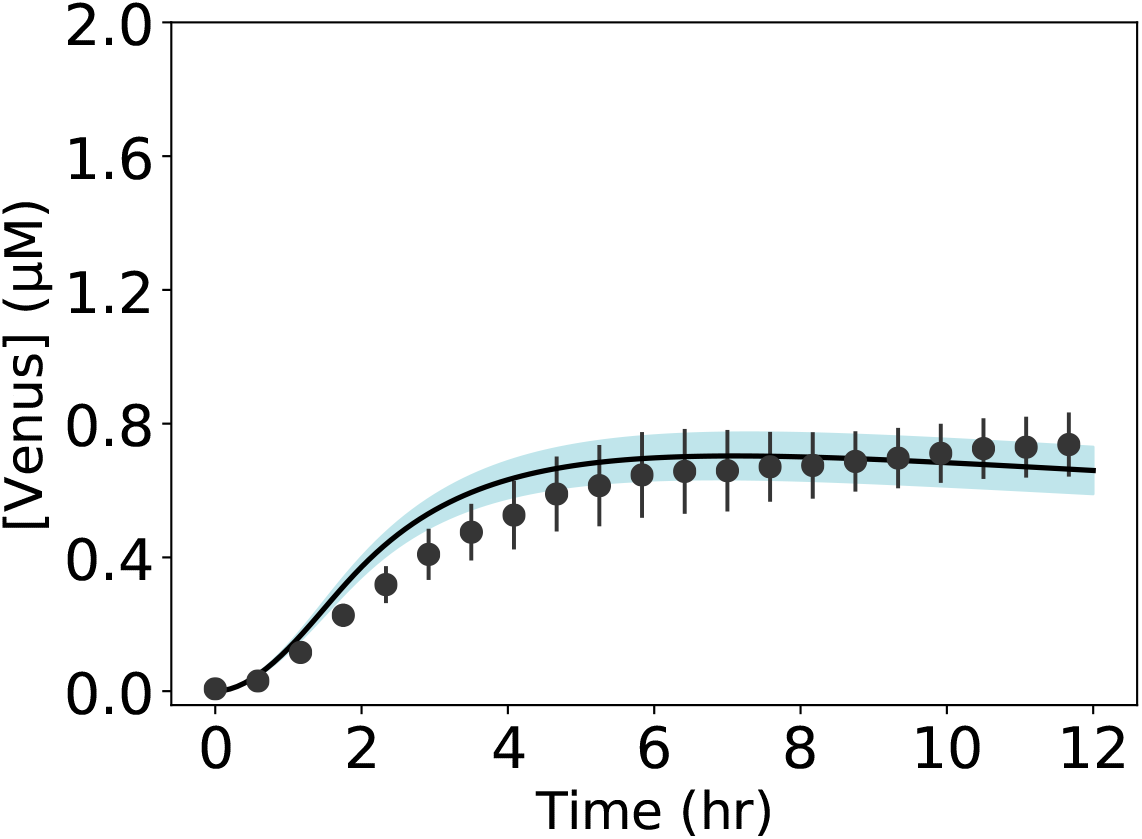
Model simulations vs. experimental measurements for the de-repressed case using 0.1 mM D-gluconate. Linear DNA used were: mP70-Venus (7 nM), P70-GntR (10 nM). Black circles represent the experimental data with the standard errors captured by the error bars across at least three replicates. JuPO-ETS parameter ensemble (N= 216) was used to calculate the mean (black line) and the 95% confidence estimate of the model simulation (blue region).

**Fig. S8:**
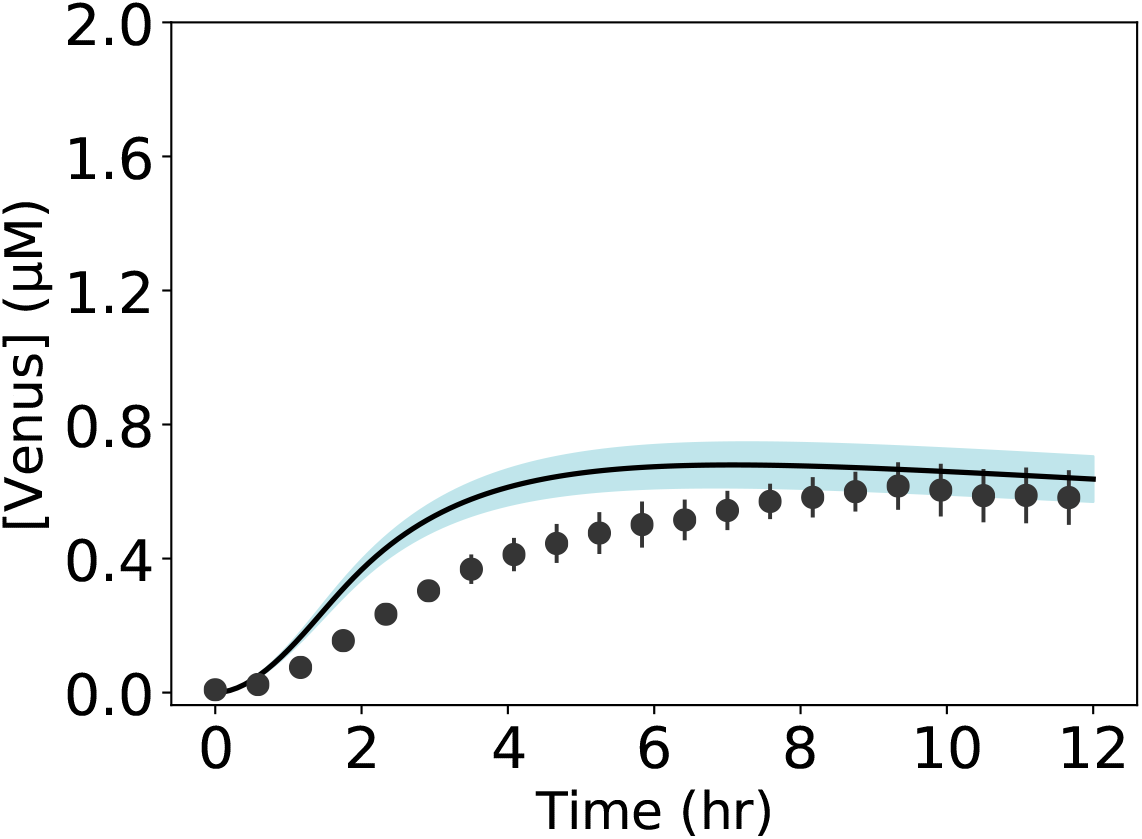
Model simulations vs. experimental measurements for the de-repressed case using 0.01 mM D-gluconate. Linear DNA used were: mP70-Venus (7 nM), P70-GntR (10 nM). Black circles represent the experimental data with the standard errors captured by the error bars across at least three replicates. JuPO-ETS parameter ensemble (N= 216) was used to calculate the mean (black line) and the 95% confidence estimate of the model simulation (blue region).

**Fig. S9:**
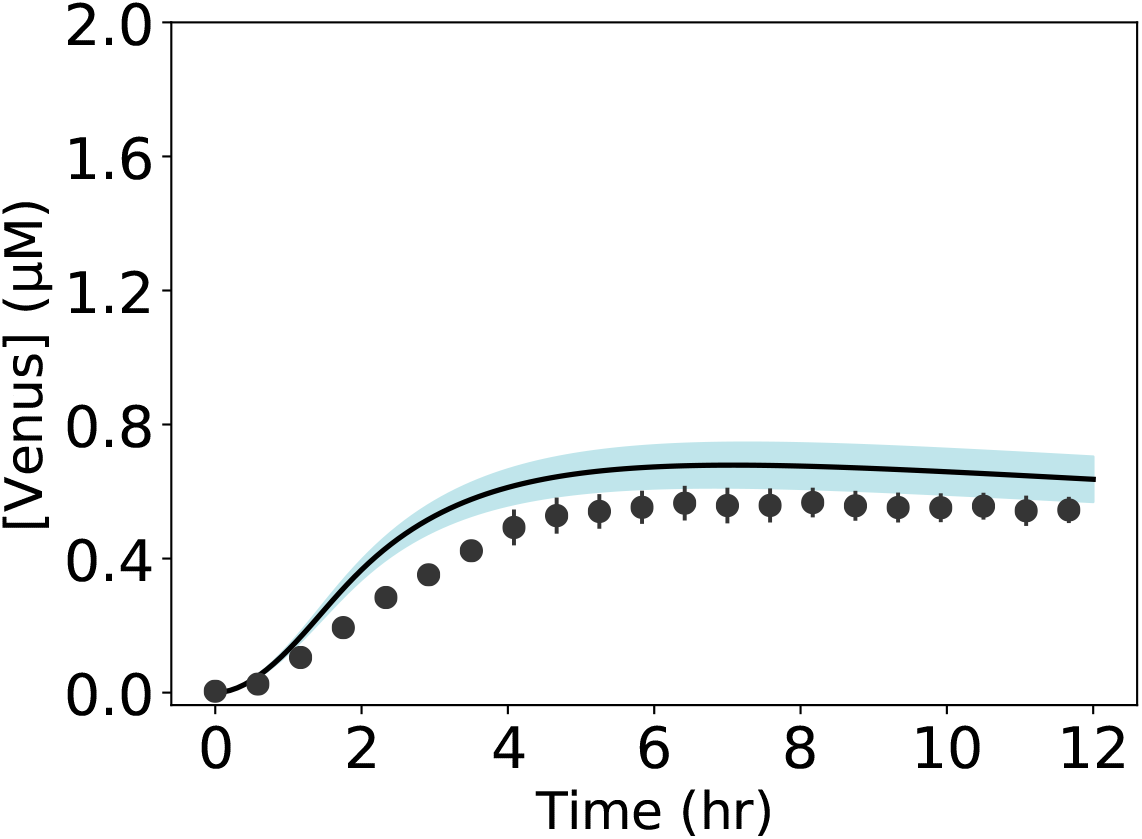
Model simulations vs. experimental measurements for the de-repressed case using 0.001 mM D-gluconate. Linear DNA used were: mP70-Venus (7 nM), P70-GntR (10 nM). Black circles represent the experimental data with the standard errors captured by the error bars across at least three replicates. JuPO-ETS parameter ensemble (N= 216) was used to calculate the mean (black line) and the 95% confidence estimate of the model simulation (blue region).

**Fig. S10:**
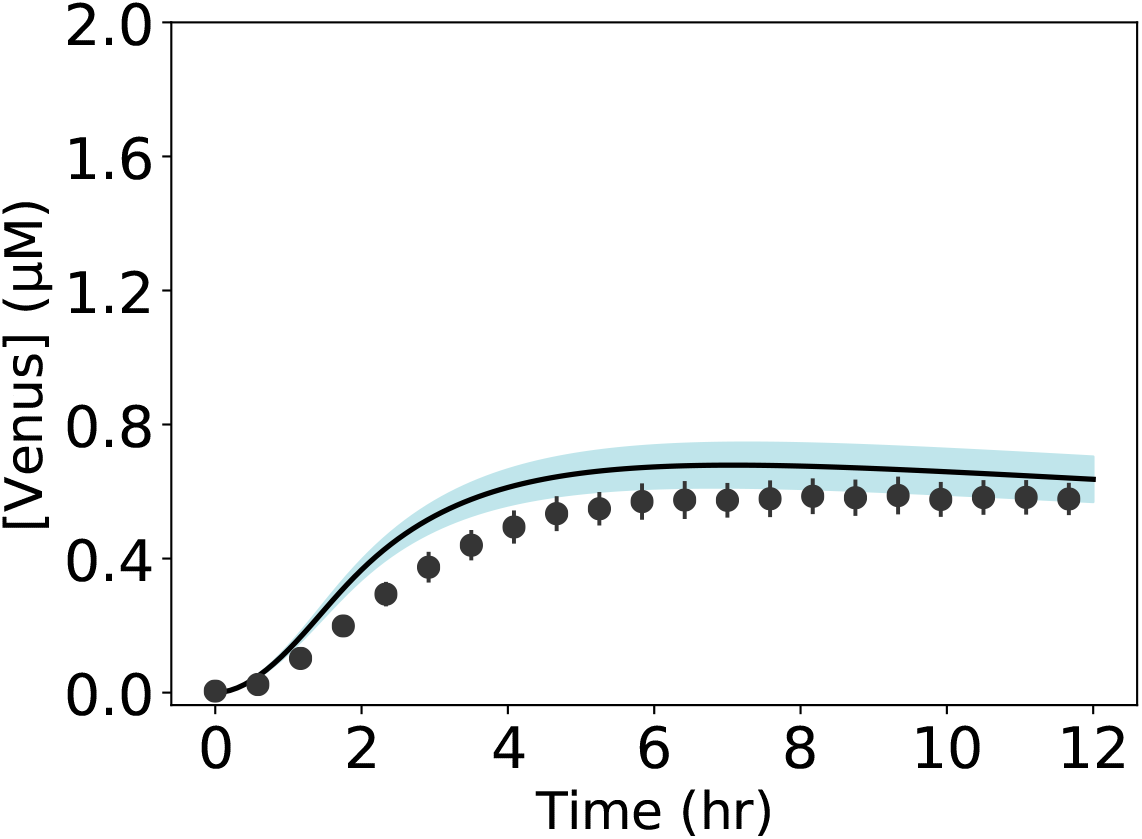
Model simulations vs. experimental measurements for the de-repressed case using 0.0001 mM D-gluconate. Linear DNA used were: mP70-Venus (7 nM), P70-GntR (10 nM). Black circles represent the experimental data with the standard errors captured by the error bars across at least three replicates. JuPO-ETS parameter ensemble (N= 216) was used to calculate the mean (black line) and the 95% confidence estimate of the model simulation (blue region).

## References

1. Swartz JR (2018) Expanding biological applications using cell-free metabolic engineering: an overview. Metabolic engineering 50: 156–172.

2. Silverman AD, Karim AS, Jewett MC (2020) Cell-free gene expression: an expanded repertoire of applications. Nature reviews genetics 21: 151–170.

3. Bundy BC, Hunt JP, Jewett MC, Swartz JR, Wood DW, et al. (2018) Cell-free biomanufacturing. Current opinion in chemical engineering 22: 177–183.

4. Vilkhovoy M, Adhikari A, Vadhin S, Varner JD (2020) The evolution of cell free biomanufacturing. Processes 8: 675.

5. Garenne D, Haines MC, Romantseva EF, Freemont P, Strychalski EA, et al. (2021) Cell-free gene expression. Nature reviews methods primers 1: 1–18.

6. Soltani M, Davis BR, Ford H, Nelson JAD, Bundy BC (2018) Reengineering cell-free protein synthesis as a biosensor: Biosensing with transcription, translation, and protein-folding. Biochemical Engineering Journal 138: 165–171.

7. Gräwe A, Dreyer A, Vornholt T, Barteczko U, Buchholz L, et al. (2019) A paper-based, cell-free biosensor system for the detection of heavy metals and date rape drugs. PloS one 14: e0210940.

8. Voyvodic PL, Bonnet J (2020) Cell-free biosensors for biomedical applications. Current opinion in biomedical engineering 13: 9–15.

9. Jung JK, Alam KK, Verosloff MS, Capdevila DA, Desmau M, et al. (2020) Cell-free biosensors for rapid detection of water contaminants. Nature biotechnology 38: 1451–1459.

10. Voyvodic PL, Pandi A, Koch M, Conejero I, Valjent E, et al. (2019) Plug-and-play metabolic transducers expand the chemical detection space of cell-free biosensors. Nature communications 10: 1–8.

11. Miwa Y, Fujita Y (1988) Purification and characterization of a repressor for the Bacillus subtilis gnt operon. The Journal of biological chemistry 263: 13252–7.

12. Fujita Y, Miwa Y (1989) Identification of an operator sequence for the Bacillus subtilis gnt operon. The journal of biological chemistry 264: 4201–6.

13. Peekhaus N, Conway T (1998) Positive and negative transcriptional regulation of the Escherichia coli gluconate gegulon gene gntT by GntR and the cyclic AMP (cAMP)-cAMP receptor protein complex. Journal of bacteriology 180: 1777–1785.

14. Yoshida KI, Miwa Y, Ohmori H, Fujita Y (1995) Analysis of an insertional operator mutation (gntOi) that affects the expression level of theBacillus subtilis gnt operon, and characterization ofgntOi suppressor mutations. Molecular and general genetics MGG 248: 583–591.

15. Daddaoua A, Corral-Lugo A, Ramos JL, Krell T (2017) Identification of gntr as regulator of the glucose metabolism in pseudomonas aeruginosa. Environmental micro-biology 19: 3721–3733.

16. Shin J, Noireaux V (2010) Efficient cell-free expression with the endogenous e. coli rna polymerase and sigma factor 70. Journal of biological engineering 4: 1–9.

17. Izu H, Adachi O, Yamada M (1997) Gene organization and transcriptional regulation of the gntrku operon involved in gluconate uptake and catabolism of escherichia coli. Journal of molecular biology 267: 778–793.

18. Adhikari A, Vilkhovoy M, Vadhin S, Lim HE, Varner JD (2020) Effective biophysical modeling of cell free transcription and translation processes. Frontiers in bioengineering and biotechnology: 1353.

19. McClure WR (1980) Rate-limiting steps in rna chain initiation. Proceedings of the National Academy of Sciences 77: 5634–5638.

20. Moon TS, Lou C, Tamsir A, Stanton BC, Voigt CA (2012) Genetic programs constructed from layered logic gates in single cells. Nature 491: 249–53.

21. Bauer J, Groneberg DA (2016) Measuring spatial accessibility of health care providers–introduction of a variable distance decay function within the floating catchment area (fca) method. PloS one 11: e0159148.

22. Stögbauer T, Windhager L, Zimmer R, Rädler JO (2012) Experiment and mathematical modeling of gene expression dynamics in a cell-free system. Integrative biology 4: 494–501.

23. Kim DM, Swartz JR (2000) Prolonging cell-free protein synthesis by selective reagent additions. Biotechnology progress 16: 385–390.

24. Whittaker JW (2013) Cell-free protein synthesis: the state of the art. Biotechnology letters 35: 143–152.

25. Mushegian AA (2016) A ribosomal strategy for magnesium deficiency. Science signaling 9: ec269–ec269.

26. Nierhaus KH (2014) Mg2+, k+, and the ribosome. Journal of bacteriology 196: 3817–3819.

27. Bassen DM, Vilkhovoy M, Minot M, Butcher JT, Varner JD (2017) Jupoets: a constrained multiobjective optimization approach to estimate biochemical model ensembles in the julia programming language. BMC systems biology 11: 1–11.

28. Rackauckas C, Nie Q (2017) Differentialequations.jl–a performant and feature-rich ecosystem for solving differential equations in julia. Journal of Open Research Software 5.

29. Morris MD (1991) Factorial sampling plans for preliminary computational experiments. Technometrics 33: 161–174.

30. Dixit VK, Rackauckas C (2022) Globalsensitivity. jl: Performant and parallel global sensitivity analysis with julia. Journal of Open Source Software 7: 4561.

31. Cuthbertson L, Nodwell JR (2013) The tetr family of regulators. Microbiology and molecular biology reviews: MMBR 77: 440–75.

32. Dimas RP, Jordan BR, Jiang XL, Martini C, Glavy JS, et al. (2019) Engineering dna recognition and allosteric response properties of tetr family proteins by using a module-swapping strategy. Nucleic acids research 47: 8913–8925.

33. Suvorova IA, Korostelev YD, Gelfand MS (2015) Gntr family of bacterial transcription factors and their dna binding motifs: structure, positioning and co-evolution. PloS one 10: e0132618.

34. Cherney LT, Cherney MM, Garen CR, Lu GJ, James MNG (2008) Crystal structure of the arginine repressor protein in complex with the dna operator from mycobacterium tuberculosis. Journal of molecular biology 384: 1330–40.

35. Lentini R, Forlin M, Martini L, Del Bianco C, Spencer AC, et al. (2013) Fluorescent proteins and in vitro genetic organization for cell-free synthetic biology. ACS synthetic biology 2: 482–489.

36. Niederholtmeyer H, Xu L, Maerkl SJ (2013) Real-time mrna measurement during an in vitro transcription and translation reaction using binary probes. ACS synthetic biology 2: 411–417.

37. Bintu L, Buchler NE, Garcia HG, Gerland U, Hwa T, et al. (2005) Transcriptional regulation by the numbers: models. Current opinion in genetics & development 15: 116–124.

38. Garamella J, Marshall R, Rustad M, Noireaux V (2016) The all e. coli tx-tl toolbox 2.0: a platform for cell-free synthetic biology. ACS synthetic biology 5: 344–355.

39. Kassavetis G, Chamberlin M (1981) Pausing and termination of transcription within the early region of bacteriophage t7 dna in vitro. Journal of Biological Chemistry 256: 2777–2786.

40. Underwood KA, Swartz JR, Puglisi JD (2005) Quantitative polysome analysis identifies limitations in bacterial cell-free protein synthesis. Biotech Bioeng 91: 425–35.

